# Habitat heterogeneity drives scale-dependent biodiversity loss in a temperate marine ecosystem

**DOI:** 10.1101/337972

**Authors:** Samuel Starko, Lauren Bailey, Elandra Creviston, Katelyn James, Alison Warren, Christopher J. Neufeld

**Author notes:** Equal contribution (Samuel Starko, Christopher J. Neufeld).

## Abstract

Biodiversity loss is driven by interacting factors operating at different spatial scales. Yet, there remains uncertainty as to how fine-scale environmental conditions mediate biological responses to broad-scale stressors. We surveyed mid-latitude kelp bed habitats to determine whether local habitat heterogeneity has mediated changes in community diversity after more than two decades of extreme temperature events, most notably the 2013-2016 heat wave. Local wave exposure conditions were key in determining responses, with some habitats remaining stable and others experiencing near complete diversity loss, leading to local declines without regional extinctions. Wave-sheltered shores, which saw the largest declines, are a very common habitat type in the Northeast Pacific and may be especially sensitive to climate-related losses in kelp diversity and abundance. Our findings highlight how local gradients can interact with global drivers to facilitate diversity loss and demonstrate how incorporating differences between habitat patches can be essential to capturing scale-dependent biodiversity loss across the landscape.

## Introduction

Ongoing biodiversity loss is expected to reduce ecosystem functioning and services (Isbell et al. 2017) but uncertainty remains about the spatial scale at which to investigate the environmental drivers of such loss (Thomas 2013, Gonzalez et al. 2016, Cardinale et al. 2018). Global stressors can interact with local factors to exacerbate or ameliorate community responses to ongoing global change (Helmuth et al. 2002, 2014, Russell and Connell 2012). Yet, fine-scale microclimatic differences between sites are often ignored by both global climate models–which predict systematic shifts in the latitudinal ranges of species (Parmesan and Yohe 2003)–, and in meta-analyses of local diversity change–which group plots only by habitat-type (e.g. forest, marsh, grassland) or by region (e.g. Vellend et al. 2013, Dornelas et al. 2014, Krumhansl et al. 2016). These common approaches, although insightful, may miss functionally important trends in community diversity change if the stresses associated with a habitat patch depend more on local conditions than on regional patterns (Tewksbury et al. 2008), or if even the most consistent declines in diversity are occurring at only a subset of sites within each habitat type. Understanding how to look for functionally-relevant biodiversity changes will therefore depend on determining the relative importance of both broad-scale and fine-scale stressors in driving community shifts through time. While much work has focussed on how broad-scale stressors are driving the biological responses of communities (Parmesan and Yohe 2003, Vellend et al. 2013, Helmuth et al. 2014, Wernberg et al. 2016), few studies have examined the role that local, fine-scale conditions play in facilitating or ameliorating them (Helmuth et al. 2014).

The rocky intertidal zone is an excellent model system for assessing the impacts of climate change on biodiversity loss (Harley 2011). Intertidal organisms live near their physiological limits (Helmuth et al. 2002) and are sensitive to air temperatures, which tend to be more variable and extreme than seawater temperatures (Helmuth et al. 2002, 2016). Local environmental gradients also have profound effects on intertidal systems. In particular, wave action plays a fundamental role in structuring intertidal communities (Leigh et al. 1987, Denny 2006). Water movement from waves can increase productivity by eliminating nutrient-depleted or oxygen-rich boundary layers that are associated with low-flow environments (Hurd 2017). Furthermore, wave splash can ameliorate the stressors associated with aerial exposure, such as desiccation and thermal stress (Helmuth et al. 2002, 2016). Given the importance of wave action on the physiology and ecology of organisms that live along rocky shorelines, exposure to waves is likely to affect how these organisms respond to global change. While thermal profiles of intertidal sites support the prediction that local conditions, such as wave exposure, will mediate community responses to climate change (Fitzhenry et al. 2004, Helmuth et al. 2016), the scarcity of appropriate baseline community data have made this difficult to test in the field.

Here we explicitly test this hypothesis by investigating the role of a local wave exposure gradient on temporal changes in intertidal kelp bed habitats in Barkley Sound, British Columbia, Canada following 22 years of broad-scale climate warming and extreme temperature events (Figs. S1-S3) (Bond et al. 2015, Levine and McPhaden 2016). Kelp communities provide many important ecosystem services to humans through rapid growth and formation of habitat for commercially harvested animals (Steneck et al. 2002), but as cold water species, kelps are expected to be sensitive to climate change. Tropicalization has already caused negative impacts on kelp forests near their range edges (Wernberg et al. 2016). However, interactions between global, regional and local processes have led to complex responses of kelp communities with large variability in the magnitude and direction of change (Wernberg et al. 2011, Krumhansl et al. 2016, Filbee-Dexter and Wernberg 2018). Studies of local-scale temporal change in the abundance of kelp and other large brown algae are increasingly common (e.g. Bennett et al. 2015, Krumhansl et al. 2016, Reed et al. 2016, Pfister et al. 2017) and collectively demonstrate that local conditions can interact with global stressors to drive ecosystem responses to change. However, these studies have focused on a small number of species and have not examined temporal changes in kelp assemblage diversity. Moreover, it still remains unclear how natural variation in site-level environmental conditions will influence the responses of kelp-dominated ecosystems to global-scale climate change or extreme climate events.

To assess temporal changes in kelp diversity, we resurveyed intertidal sites (n=49) in 2017 that had previously been surveyed by Druehl & Elliot between 1993 and 1995 (Druehl, L.D. and Elliot, C.T.J. 1996). We identified all kelp species present at each site and quantified abundance at a majority of sites (n = 45). Surveys (performed between June 20 and Sept 9, 2017) were conducted on 20-50 m stretches of coastline and included the entire intertidal region, from Lower Low Water Large Tide (LLWLT) to the upper limit of marine organisms. A subset of these sites (N = 17) were surveyed in both June and September and no differences in kelp diversity were detected during this time. Sites occurred broadly throughout the region, and were situated across a range of wave exposures, slopes, aspects, and types of rocky substrates. Public data from nearby lighthouses show that both sea surface (SST) and air temperatures in Barkley Sound have reached abnormal highs between 1995 and 2017 with especially high temperatures occurring during the 2014-2016 heatwave (Figs S1-S3, Supplemental Discussion). Moreover, both air temperatures and water temperatures were higher between the 5-year period of 2013 – 2017 than between 1991 – 1995 (Fig S1).

## Methods

### Study system

Barkley Sound, on the southwest coast of Vancouver Island, Canada, is a nearly 30 km wide inlet containing hundreds of islands. As such, it provides a wide range of local microhabitats. Both wave-sheltered and wave-exposed sites are located throughout the area both near and far from the opening to the sound. Sites were accessed by boat. Historical survey data spans 1993-1995 with most (n = 46) sites sampled twice in 1993 (n = 19) or 1994 (n = 27) and 1995. However, three sites were only sampled in either 1993 (n = 2) or 1994 (n = 1) and not in 1995 (Supplementary Table 1). Sites were located using GPS coordinates, photographs, and descriptions recorded in the original surveys. In particular, most sites were located using a photo that was often annotated with the exact location of the transect. A thorough description of the site was also given by the authors and allowed for location of some sites that did not have photographs. Sites were only resurveyed if they could be definitively located in at least one of these two ways using distinct geographic landmarks.

### Survey techniques

Surveys were conducted following the methods of the original surveyors. Survey sites were uniform lengths of shoreline and included the area between the high tide line and LLWLT (approximately 3 m vertical distance). Presence and absence of all kelp species were determined for the entire survey area by carefully identifying all kelp species present in an area by morphology. Kelps are large, seasonally persistent and are easy to distinguish based on conspicuous morphological features (Druehl, L.D. and Elliot, C.T.J. 1996). Thus, both our surveys and those done by the original surveyors were likely to result in unbiased, reproducible data. In order to quantify abundance, the intertidal was blocked into four zones: high intertidal (approx. > 2.5 m), mid intertidal (approx. 1.2 – 2.5 m), low intertidal (approx. 0.2 – 1.2 m) and shallow subtidal (0 – 0.2 m). Abundance of each species, in each zone, was then quantified based on visual estimation of percentage cover categories: absent (0 %), rare (< 5 %), common (6 – 20 %) and abundant (21 – 100 %). A species’ assigned abundance was then taken from the zone of its greatest abundance.

### Wave exposure quantification

Quantifying wave exposure is a known challenge to intertidal biologists, as local topography can influence water velocities in ways that many geographical indices fail to capture (Helmuth and Denny 2003). For characterization of sites, we used the site-specific wave-exposure categories provided by the original Barkley Sound surveyors (Druehl, L.D. and Elliot, C.T.J. 1996). This method categorized sites based on direct, qualitative observations of water motion (Topinka et al. 2009) and the presence of indicator species. We used a cartographical method previously developed (Burns and Neufeld 2009) and tested (Neufeld et al. 2017) in Barkley Sound to further test the validity of these categories. In brief, this method is a continuous index derived from the angle of unimpeded exposure to the predominant direction of offshore swell (southwest). It is therefore only effective for SW facing sites (Neufeld et al. 2017). We used all of our SW (180-270°) facing sites (N = 26) to ground-truth these wave exposure categories (ANOVA on ranks: F = 17.24, P < 0.001; Fig S6) and showed that sites categorized as “Protected” had significantly lower wave exposure index measures than “Moderate” and “Exposed” (Tukey HSD: P < 0.001 for both). There was a near significant trend suggesting a difference between “Moderate” and “Exposed” sites (Tukey HSD: P = 0.0637). Together, this suggests that our wave exposure categories were appropriate.

### Shoreline classification

In order to determine how any wave exposure-specific responses might scale up across the landscape, we examined the distribution of rocky habitats of different wave exposures across the North American Pacific coast using a comprehensive georeferenced linear shoreline dataset called ShoreZone (Howes 2001). ShoreZone data spans from Oregon to Alaska and is based on expert classification of shoreline units using low-elevation aerial imagery obtained from fixed-wing aircraft or helicopter and relevant geographic features. During segment classification, each shoreline unit is assigned a substrate class from high-resolution imagery, and assigned a wave-exposure class using a combination of fetch calculations and geographic and biotic features. For the current study, predominantly rocky shoreline was identified by selecting all shoreline units from this dataset which contained at least 25% rocky substrate (Shorezone coastal classes 1-20). Regional totals of the extent of shoreline containing only bedrock (ShoreZone coastal classes 1-5) were 44-78% shorter than regional totals from mostly rocky shoreline but produced the same patterns of relative habitat types between regions. Because the average shoreline unit in the ShoreZone dataset is between 300m and 500m long (much longer than our 20-50 m surveys), the ShoreZone wave exposure classification is not able to resolve small scale differences in exposure that fall within a single shoreline unit. Importantly, Shorezone produced similar categorizations as Druehl & Elliot: 96% of sites were within one wave exposure category of one another (and classifications at 67% of sites agreed completely). Sites that differed between methods included a tidepool that was set back from the shore and protected from incoming waves, three exposed headland sites in areas that were otherwise largely sheltered from waves, and two sites that were located on the wave-sheltered side of islands that were near the mouth of the sound where overall wave exposure is greater. Barring these few exceptions that arose largely due to differences in the scale at which wave exposure was assessed, the overall concordance of the two independent approaches suggests that scaling up to the broader region using the ShoreZone dataset is appropriate.

### Statistical analyses

All analyses were performed in R 3.4.1 (R Development Core Team, 2008). To assess whether wave exposure and survey year significantly influenced the richness and abundance of kelps at each site, we used ANOVAs on site-level proportional responses (ratio of historic to modern values) with wave exposure (fixed factor, three levels) as an explanatory variable. Comparisons were made to averages of historical survey data when more than one year was sampled during the 1993-1995 surveys (n = 46). Rarefaction and regional species pool extrapolation were performed using the “vegan” package (Oksanen et al. 2018) in R. Modern (2017) trends were compared to 1995 data when available (n = 46) or else 1993 (n = 2) or 1994 (n = 1) data were used. However, separate comparisons to 1993-1994 (n = 49) and 1995 (n = 46) are provided in the supplementary (Fig S7, S8). While diversity of annual species was generally lowest in 1995, comparing separately to the different initial survey years did not influence our overall results. To rule out spatial effects, we tested for spatial correlation of proportional richness responses using Moran’s I in the R package “ape”.

## Results & Discussion

Across the study system, we found widespread declines in kelp species richness and abundance mediated by local variation in wave action. Kelp species richness has not changed significantly at wave-exposed sites (Paired t-test: t = 0.63511, df = 9, p = 0.5412), while kelp communities at wave sheltered sites have been reduced to between zero and three species, regardless of their historical diversity (Fig. 1, Fig. 2). This is exemplified by a significant effect of wave exposure on proportional change in richness (ANOVA: F_47,1_ =17.27, P=0.00014, Fig 2A, Fig S4). This habitat-dependent decline in diversity is further supported by the shallow slope of the rarefaction curve for wave-sheltered and moderate sites in 2017 (Fig 3D). At all wave exposures, average abundance also declined and this decline was greatest at wave sheltered sites (Fig 2B; ANOVA: F_43,1_ = 4.396, P=0.042).

**Figure 1.**
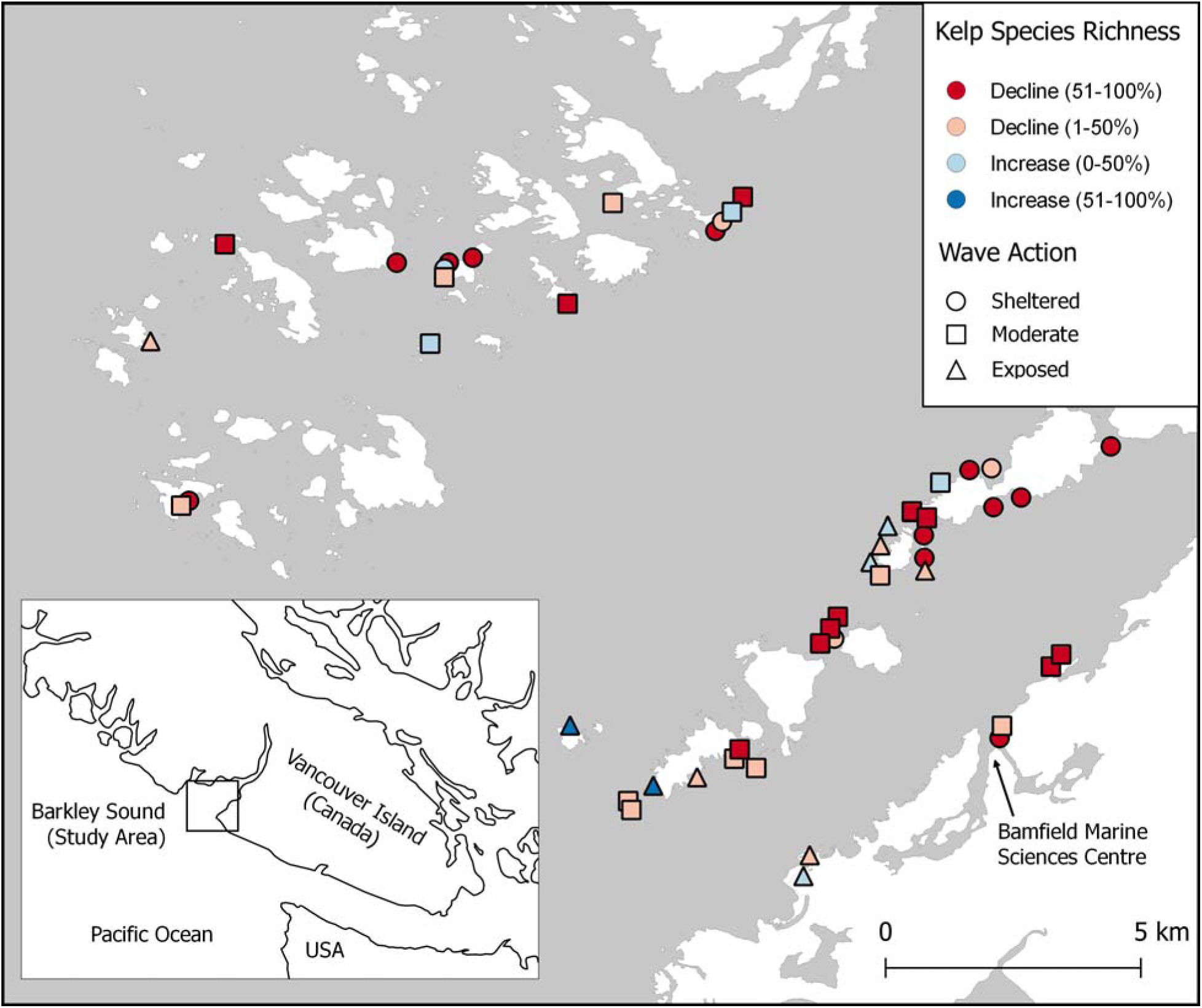
Study region. Study sites (n = 49) are coded for magnitude of change in kelp species richness (colour) and relative exposure to waves (shape). Due to the close proximity of some sites, some symbols have been moved slightly to avoid obscuring overlapping symbols. There was no effect of the spatial distribution of sites on proportional kelp richness change (Moran’s I: I = 0.0303, p = 0.49676).

**Figure 2.**
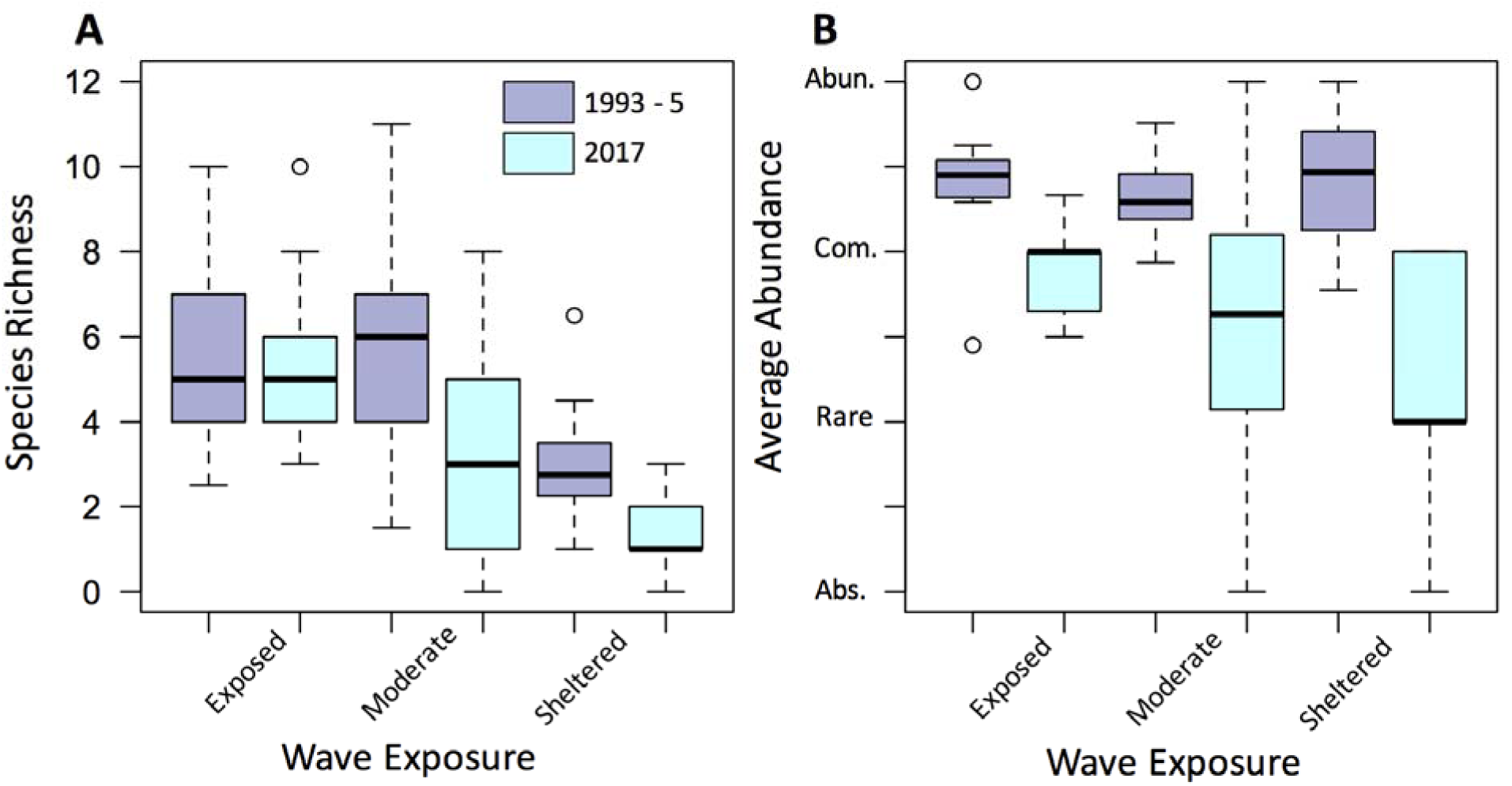
(A) Richness and (B) average abundance of species at each site (absent = 0, rare = 1, common = 2, abundant = 3) for historical (average of 1993 – 1995) and modern (2017) observations. There is a significant effect of wave exposure on site-wise proportional changes in richness (ANOVA: F_47,1_ =17.27, P=0.000136) and abundance (ANOVA: F_43,1_ = 4.396, P=0.0420).

**Figure 3.**
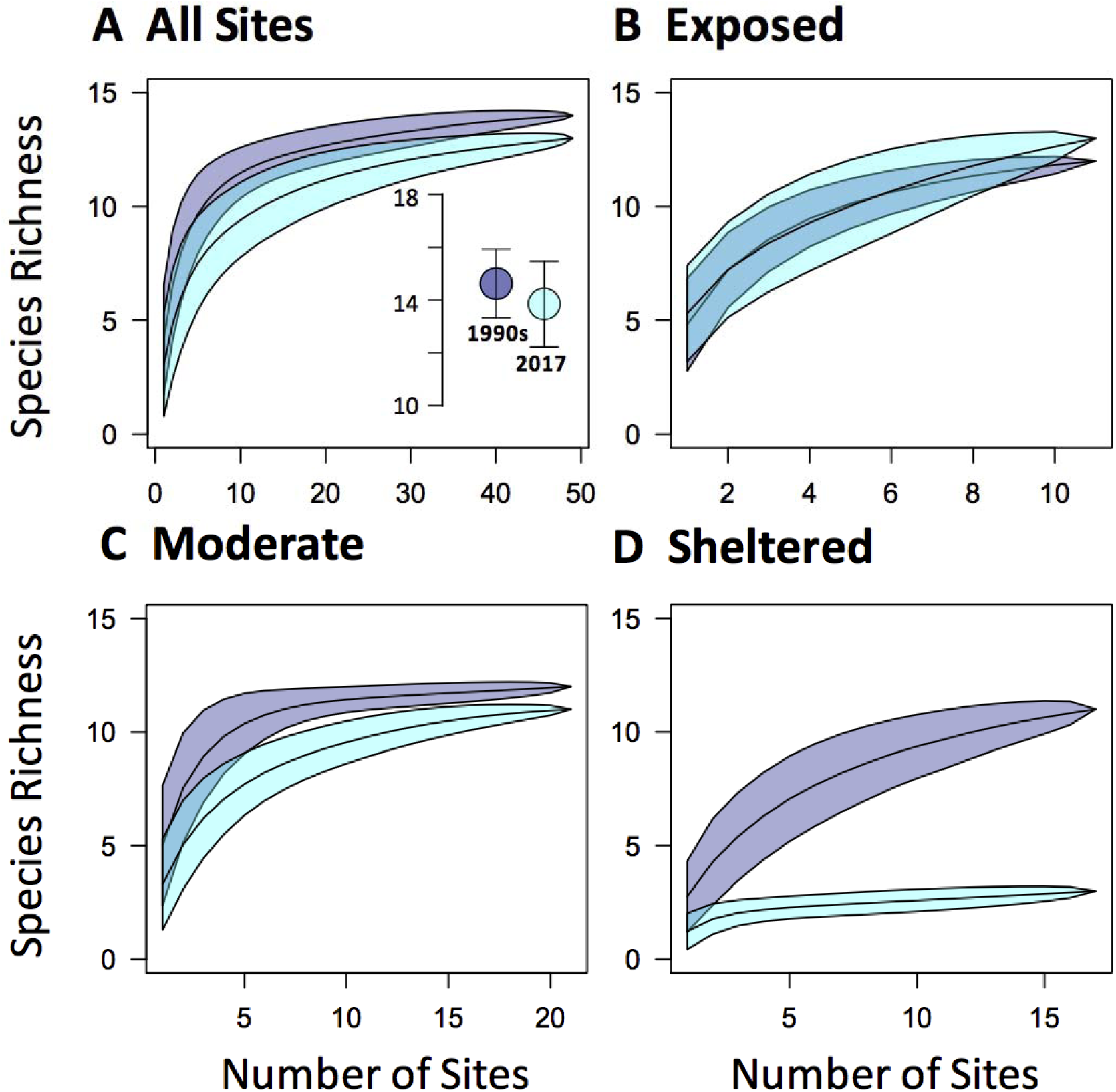
Rarefaction curves (10,000 iterations) for modern (2017) and historic (1995 for 46 sites, 1993 for 2 sites and 1994 for 1 site) surveys of sites in Barkley Sound. Analyses are broken down by (A) all sites, and then for (B) wave exposed, (C) moderately exposed sites and (D) sheltered sites plotted separately. The inset in (A) shows the results of a bootstrap extrapolation of total regional species richness of the Sound.

Changes of between-site diversity (i.e. beta and gamma) also depended largely on wave exposure. While rarefaction curves of the entire region are similar between years across all sites, sheltered sites saw marked declines in both slope and asymptote (Fig 3), indicating spatial homogenization of these communities (Tyler and Kowalewski 2017). Moreover, while at exposed sites the total number of species detected was slightly larger than in 1995, the total species pool at sheltered sites declined from 11 species to 3 (Fig 2) with one of those species only present at one site. Despite these widespread declines in the number of species found at sheltered sites, species richness across the region did not change (Fig 3A inset, Fig S5; Supplementary Discussion). Taken together, our results show a strong disconnect in how kelp diversity has changed across scales and habitats: richness of the regional kelp assemblage has not changed, while local richness and average abundance has declined markedly, with diversity loss concentrated on wave-sheltered shores.

It is unclear whether we have documented a permanent loss of kelp habitat or a temporary response to recent oceanographic events (Lorenzo and Mantua 2016). Yet, the magnitude of the species loss and the multiannual life cycle of many affected kelp species, greatly suggest that that our results represent widespread declines in local kelp diversity across Barkley Sound. Analysis of data from nearby lighthouses shows no clear, linear increase in either air or water temperature over the 22 year period but show an obvious signal of anomalous temperatures during the recent 2013-2016 heat wave (Fig S2,S3). Heatwaves have been shown to have large-scale impacts on subtidal kelps (Filbee-Dexter and Wernberg 2018) but this has not been previously documented in intertidal kelp communities. Regardless of the timescale over which these declines occurred, our results suggest that wave-sheltered habitats are more sensitive to regional stressors than wave exposed habitats. Splashing of cool water at exposed sites could alleviate air temperature stress during low tide, leading to the patterns that we show here (Harley and Helmuth 2003) or local mixing at sites with increased water motion could mediate these stresses by preventing pockets of warm water from forming at small scales. During the recent 2014 – 2015 “Blob” and the 2016 El Nino, nitrogen levels were also abnormally low (Lorenzo and Mantua 2016), (Reed et al. 2016). Nutrient availability may influence thermal tolerance and so multiple stressors could contribute to these declines (Thomas et al. 2017).

A few key implications arise from these findings. First, given the important ecological role of kelp (Duggins et al. 1989, Steneck et al. 2002, Teagle et al. 2017), the declines that we document are likely to have cascading effects on the diversity of other organisms and on ecosystem functioning of intertidal communities (Teagle et al. 2017). While the affected kelp communities may yet recover following the 2013-2016 heatwave, our results suggest that kelp communities at wave-sheltered sites may be particularly sensitive to increasing global temperatures and heat wave frequency. It could be hypothesized that declines on wave-sheltered shores may not affect regional productivity or habitat availability as much as would declines on wave-exposed shorelines, which are more diverse and more productive (Leigh et al. 1987). Yet, positive interactions generated by kelp canopies may be especially important on wave-sheltered shores because these shores are more physiologically stressful (Bruno et al. 2003). Furthermore, the lower diversity and productivity of sheltered shorelines is far outweighed by their sheer abundance in the Northeast Pacific (Fig. 4). We show that 57,000 km of wave-sheltered rocky shoreline exists from Oregon to central Alaska, virtually all of which (99.8%) occurs north (and east) of Washington’s outer coast (Fig. 4A). Therefore, even small changes in kelp diversity on more sensitive wave-sheltered shores could have large effects on intertidal productivity if magnified across the landscape. While some of this shoreline may not suitable kelp habitat due to limitations from salinity and other factors, it is clear from our analyses that wave sheltered shorelines are common in northern Washington, British Columbia and Alaska. Given that these types of habitats are uncommon further south, it is possible that some northern shorelines will experience losses before southern ones.

**Figure 4.**
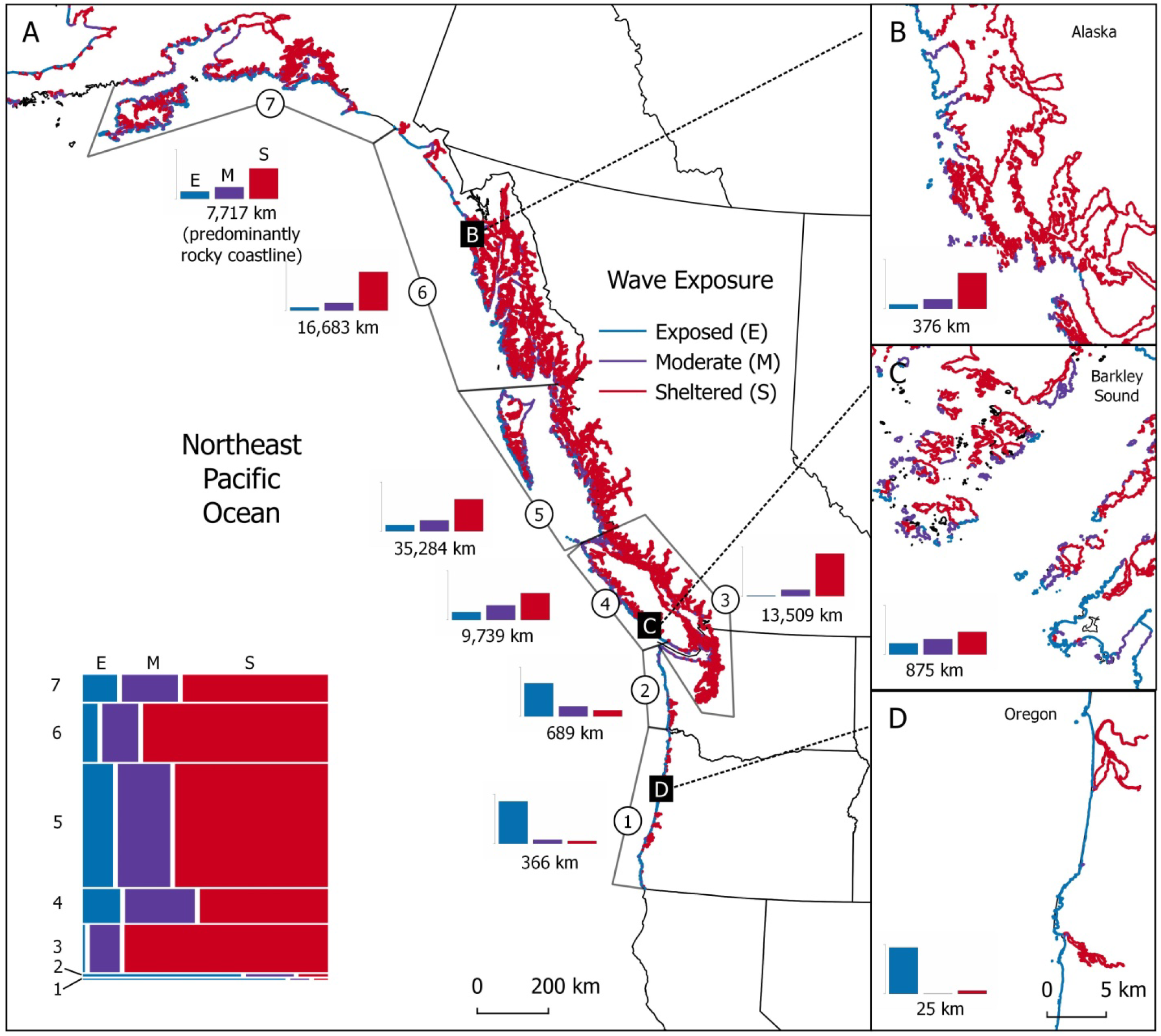
Northeast Pacific intertidal habitat classified by wave exposure. Wave-sheltered habitat makes up the majority of Northeast Pacific shorelines (A) and is abundant in British Columbia (C) and Alaska (B) but rare along the outer coast of Washington and Oregon (D). Bar plots show the proportions of rocky shoreline of different wave-exposures and are accompanied by the length of predominantly rocky shoreline in each region (A) or in each inset (B-D). Mosaic plot inset in A shows the relative proportions of rocky shoreline of different wave-exposures scaled by the length of coastline for each region. Regions from South to North: (1) Oregon coast, (2) Washington outer coast, (3) Salish Sea, Puget Sound, and Strait of Georgia, (4) western Vancouver Island, (5) northern British Columbia, (6) southern Alaska, (7) central Alaska.

The apparent resilience of wave exposed kelp beds also has important implications for conservation and management. Recent efforts to understand biodiversity change in ecologically important biogenic habitats have identified areas where ecosystems are performing substantially better (“bright spots”) or worse (“dark spots”) than average (Cinner et al. 2016). Our results demonstrate that a fine-scale environmental gradient, that can vary over tens of metres (Helmuth and Denny 2003), has mediated the responses of kelp. Thus, kelp bright spots and dark spots may be located close together in space and should be examined at this fine scale. Wave exposed sites are highly productive and often represent hotspots of diversity in our system (Druehl, L.D. and Elliot, C.T.J. 1996). Our results demonstrate that these sites may also be especially resilient against ongoing global change. Wave exposed sites may act as refugia during times of extreme climate stress, potentially buffering kelp ecosystems against regional extinctions and playing a key role in maintaining species diversity. The role of climatic refuges in maintaining species diversity through geological time has been widely discussed in the paleoecological literature (Haffer 1969, e.g. Willis and Whittaker 2000). However, few studies illustrate this phenomenon under current climate change. Given the distribution of wave exposure in the Northeast Pacific (Fig 4), dark spots are likely to be common. Although wave exposed sites might maintain regional diversity, abundant dark spots could have profound effects on ecosystem functioning and services (Loreau et al. 2001).

In addition to reductions in diversity, we document widespread declines in the abundance of intertidal kelps in Barkley Sound. While the magnitude of decline was dependent on wave exposure (Fig 2B) and varied between species (Fig S9-S10), sites from all wave exposure categories declined significantly in average kelp abundance. Losses of kelp cover are common and a recent global meta-analysis found that more than one third of published subtidal kelp bed surveys show declines over the past 50 years – significantly more than had increased (Krumhansl et al. 2016). While many negatively affected kelp forest ecosystems are found near the warm-edge of kelps’ latitudinal range(Wernberg et al. 2011, 2016, Assis et al. 2017, Filbee-Dexter and Wernberg 2018), our data suggest that similar declines have occurred in the intertidal zones of British Columbia, reasonably far from the warmer latitudinal limit of northeast Pacific kelp ecosystems (Bolton 2010). This supports previous work suggesting that central-and not just edge-populations of brown algae may be susceptible to large-scale increases in temperature (Bennett et al. 2015).

Although declines may be attributable to recent heatwave events occurring over short timescales (Lorenzo and Mantua 2016) (Fig S2-S3), rather than a response to gradual warming, the recovery from ecosystem-wide declines may not occur rapidly in either case. Four of our sites lost all kelp species and thirteen others were reduced to a single, sometimes rare (< 5 % cover) species. Thus, many of our sites have experienced complete or near-complete collapses of kelp-dominated communities. Climate-related phase shifts from communities dominated by kelps to those dominated by other, less productive seaweed species, such as turfs, have unfavourable consequences for coastal productivity and ecosystem services (Filbee-Dexter and Wernberg 2018). Phase shifts between kelp and turf or fucoid dominated ecosystems have been occurring in various regions worldwide for decades and many have yet to recover following initial kelp bed collapse (Filbee-Dexter et al. 2016, Filbee-Dexter and Wernberg 2018). This hysteresis, driven by feedbacks between biotic factors and the physical environment, is responsible for the resilience of degraded ecosystems worldwide (Suding et al. 2004, O’Brien and Scheibling 2018).

A broader implication of our results is that important biodiversity loss could easily remain hidden from studies not specifically designed with habitat heterogeneity in mind. Declines could be concentrated in only some habitats that may be stressful and low diversity to begin with. If these marginal habitats lack novel diversity, studies that focus on regional patterns without considering habitat heterogeneity could miss losses caused by local gradients. Thus, habitat heterogeneity may directly contribute to the disconnect between diversity measurements taken at different scales (Vellend et al. 2013, Dornelas et al. 2014). Importantly, this means that without considering differences in site-level conditions, we would have missed the widespread biotic homogenization that has occurred only at wave-sheltered sites in our system. Capturing these losses in beta diversity is important to monitoring and conservation efforts because ecosystem functionality depends on having many species combinations across the landscape (Isbell et al. 2017, Fanin et al. 2017). Yet, while our results support growing evidence that local habitat heterogeneity explains important variation in diversity loss (Harley 2011, Russell and Connell 2012), few studies that examine responses to climate change incorporate these gradients into their analyses. Our results point to the need for a framework that better incorporates the interacting effects of stressors at different scales. Such an approach would hold much promise for identifying and predicting diversity loss not only at species’ range edges but also along local gradients throughout the range of each species.

Heterogeneity in environmental variables (like wave exposure) is ubiquitous in the natural world, but its importance in determining the responses of communities to climate change is often underappreciated (Helmuth et al. 2014). As the climate warms and average conditions change, heterogeneity of habitats will lead to variation in microclimates (Helmuth et al. 2002, Fitzhenry et al. 2004) and thus will affect the biological responses of organisms. Therefore, rather than assess average responses across all communities in a region or across the globe (Vellend et al. 2013, Dornelas et al. 2014), we should work to identify the habitats that are most vulnerable to declines and determine whether they are abundant enough to influence ecosystem functioning across the landscape. Consistent declines across all habitat patches or at the most diverse, high quality habitats are likely not reasonable predictions for how communities will respond to climate change (Elahi et al. 2015). Instead, diversity may be lost from marginal habitats and, if these habitats are common, the consequences to ecosystem functioning could be profound.

## Acknowledgments

This work would not have been possible without the initial surveys which were conducted by Druehl and Elliot and funded by Parks Canada. We thank the Huu-ay-aht First Nations, S. Rogers, B. Anholt, S. Gray, B. Rogers, T. Eastham, N. Wiewel and other Bamfield Marine Sciences Centre (BMSC) staff for making this work logistically feasible. Students in the 2017 Coastal Community Ecology class at BMSC contributed to field logistics and planning. We thank C. Harley, P. Thompson and M. Whalen for their many insightful comments on an earlier version of this paper, and thank J. Sunday for helpful feedback on data interpretation. SS and CJN designed the study, collected and analyzed the data, and wrote the paper. LB, EC, KJ, and AW assisted with designing the study and collecting the data, and contributed to writing and editing the paper. The authors declare no competing interests.

